# Biogenesis and antigen presentation properties of dendritic cell-derived apoptotic bodies

**DOI:** 10.64898/2026.07.04.734104

**Authors:** Amy L. Hodge, Jascinta P. Santavanond, Bo Shi, Sarah Caruso, Sara Oveissi, Caitlin Vella, Omar Audi, Dilara C. Ozkocak, Stephanie F. Rutter, Thanh Kha Phan, Lanzhou Jiang, Satoko Arakawa, Shigeomi Shimizu, Ikuyo Yoshino, Georgia K. Atkin-Smith, Gemma F. Ryan, Weisan Chen, Jieru Deng, Mark D. Hulett, Amy A. Baxter, Ivan K. H. Poon

## Abstract

Dendritic cells (DCs) are an important type of antigen presentation cell that regulate immunity by initiating antigen-specific immunity and tolerance through T cell activation. The interaction between DCs and T cells can be mediated through direct cell-cell contact or via the release of extracellular vesicles (EVs) from DCs that harbour antigen presentation machineries. Although small EVs (<200 nm in diameter) such as exosomes released by DCs have been shown to regulate immunity, whether other EV subtypes, in particular those that are released by dying DCs due to homeostatic turnover or following infection, can modulate immune responses is not defined. In this study, we demonstrated that DCs undergoing apoptosis can generate a subclass of large EVs (∼1,000-5,000 nm in diameter) known as apoptotic bodies (ApoBDs) via distinct morphological steps. Mechanistically, ApoBD formation by apoptotic DCs is regulated by Rho-associated kinase 1 and T-type calcium channels. Functionally, DC-derived ApoBDs were found to mediate direct antigen presentation. These data demonstrate a novel function of ApoBDs and highlight the ability of apoptotic materials derived from dying DCs to continue mediating intercellular communication and regulating immune responses.

## Introduction

In the human body, approximately 200-300 billion cells are turned over daily through the programmed cell death process known as apoptosis (Danial & Korsmeyer, 2004). During apoptosis, dying cells can release soluble factors such as metabolites and expose phospholipids such as phosphatidylserine (PtdSer) to communicate with neighbouring cells to coordinate rapid removal of apoptotic materials as well as promoting anti-inflammatory and wound healing responses (Nagata *et al*, 2010; Poon & Ravichandran, 2024). In addition to soluble factors, communication between apoptotic cells and neighbouring cells can also be mediated via extracellular vesicles (EVs). Apoptotic cells can undergo sequential morphological changes to drive cell fragmentation and the formation of membrane-bound EVs known as apoptotic bodies (ApoBDs) (Atkin-Smith & Poon, 2017). ApoBDs are typically described as 1-5 μm in diameter and are categorized as a subclass of large EVs that are generated exclusively during apoptosis. Notably, ApoBDs have been shown to aid intercellular communication through biomolecules that are packaged within or bound on the vesicle including DNA, RNA, signalling molecules, cytokines, chemokines as well as pathogens (Caruso & Poon, 2018). Of particular interest, ApoBDs can also harbour cell surface molecules that correspond to the parental cell including lineage-specific markers (Jiang *et al*, 2017) as well as functional signalling molecules (Ma *et al*, 2019). For example, ApoBDs derived from mature osteoclasts contain membranous RANKL on the vesicle surface that can activate RANKL reverse signalling in pre-osteoblast to promote osteogenic differentiation (Ma *et al*, 2019). Since a range of cell types including different subsets of immune cells can undergo apoptosis under homeostatic conditions for cell turnover as well as in disease settings such as during infections (Atkin-Smith *et al*, 2018; Wallach & Kang, 2018; Ekert & Vaux, 1997), ApoBDs generated under these settings can potentially mediate important intercellular communication processes. The formation of ApoBDs has been described for a number of immune cell types including thymocytes (Ohyama *et al*, 1985), T cells (Poon *et al*, 2014), B cells (Grootveld *et al*, 2023) and monocytes (Atkin-Smith *et al*, 2015). However, whether dendritic cells (DCs), a key antigen presentation cells of the immune system, can undergo cell disassembly during apoptosis to form ApoBDs and the corresponding functional significance of this process has not been investigated.

DCs are professional antigen presenting cells that are present throughout all tissues. There are two major DC subsets; conventional/classical DCs and plasmacytoid DCs, both of which are critical in sampling their environment and function to stimulate an adaptive immune response by initiating antigen-specific immunity or tolerance (Hilligan & Ronchese, 2020; Théry & Amigorena, 2001). Antigen presentation is mediated through the display of peptides loaded on Major Histocompatibility Class I (MHC-I) or MHC-II molecules, in cooperation with T cell costimulatory molecules (e.g., CD80 and CD86). Peptide-specific cognate T cell receptors on CD4^+^ or CD8^+^ T cells can recognize these peptide-loaded MHC (p-MHC) complexes and become activated, inducing their effector functions. Although the antigen presentation process and the functional roles of different DC subsets have been studied extensively (Hilligan & Ronchese, 2020; Théry & Amigorena, 2001), whether DCs can continue to influence immunity after they have died is not well defined. Since DCs undergo apoptosis under steady state conditions (Hildeman *et al*, 2007; Kamath *et al*, 2002), as well as following viral infection (Kushwah & Hu, 2010) and antigen presentation via cytotoxic CD8^+^ T cells (Yang *et al*, 2006), apoptotic materials generated from apoptotic DCs can potentially regulate immune responses through their interaction with antigen-specific T cells. In this study, we described for the first time the ability of DCs to generate ApoBDs under homeostatic and apoptosis-promoting conditions. Mechanistically, time-lapse microscopy and flow cytometry studies revealed that apoptotic DCs undergo disassembly through the formation of dynamic membrane blebs, as regulated by Rho-associated kinase 1 (ROCK1), and formation of thin membrane protrusions called apoptopodia, as regulated by T-type calcium channels. Furthermore, DC-derived ApoBDs (DC-ApoBDs) retained MHC-I as well as T cell costimulatory molecules CD80 and CD86. Functionally, utilising the ovalbumin (OVA) antigen presentation and influenza A virus (IAV) infection models, we demonstrated the ability of DC-derived ApoBDs to present antigen directly to CD8^+^ T cells. Taken together, this study describes the biogenesis and function of ApoBDs generated from apoptotic DCs, with broad implications in their role of regulating immune responses.

## Results

### Generation of DC-ApoBDs under *in vivo* and *in vitro* conditions

DCs routinely undergo apoptosis during steady state conditions to maintain homeostasis (Hildeman *et al*, 2007; Kamath *et al*, 2002). Furthermore, maintenance of DC numbers via apoptosis is imperative as DC accumulation due to reduced levels of apoptosis can lead to systemic autoimmunity (Chen *et al*, 2006, 2007). Therefore, we first asked whether we could detect the presence of apoptotic DCs and DC-derived ApoBDs under steady state conditions in thymic and spleen tissues (Figure 1A). DCs were identified based on cell surface markers CD45^+^ CD11c^+^ MHCII^+^ and a relatively small proportion of apoptotic DCs were detected in the thymus and spleen (Figure 1B), based on staining with annexin A5 (A5, indicative of PtdSer exposure) and a caspase 3/7 fluorescent inhibitor of caspase activity probe FLICA (FAM-DVED-FMK, indicative of caspase 3/7 activation). Notably, distinct DC-derived ApoBDs (FSC^low^, A5^+^, FLICA^+^) were also identified in the thymus and spleen (Figure 1C, Supplementary Figure 1A-C). DC-derived ApoBDs from the thymus and spleen exhibited intermediate level of CD11c and MHCII surface markers expression when compared to viable DCs (Figure 1D). Surface markers for other cell types such as T cells, B cells and neutrophils (CD3, B220 and Ly6G, respectively), are not readily detected on DC-derived ApoBDs (Supplementary Figure 1B and D). Next, we sought to determine the presence of DC-derived ApoBDs upon induction of apoptosis *in vivo* by X-ray irradiation (Figure 1E). Whole-body irradiation treatment resulted in a decrease in the number of viable thymic DCs and an increase in apoptotic DCs when compared to the steady state condition (Figure 1F). Furthermore, an increase in the number of DC-derived ApoBDs were also detected in the thymus following irradiation treatment (Figure 1G). Altogether, these data demonstrate the presence of DC-derived ApoBDs in the thymus and spleen during steady state and their levels increase upon apoptosis induction.

**Figure 1.**
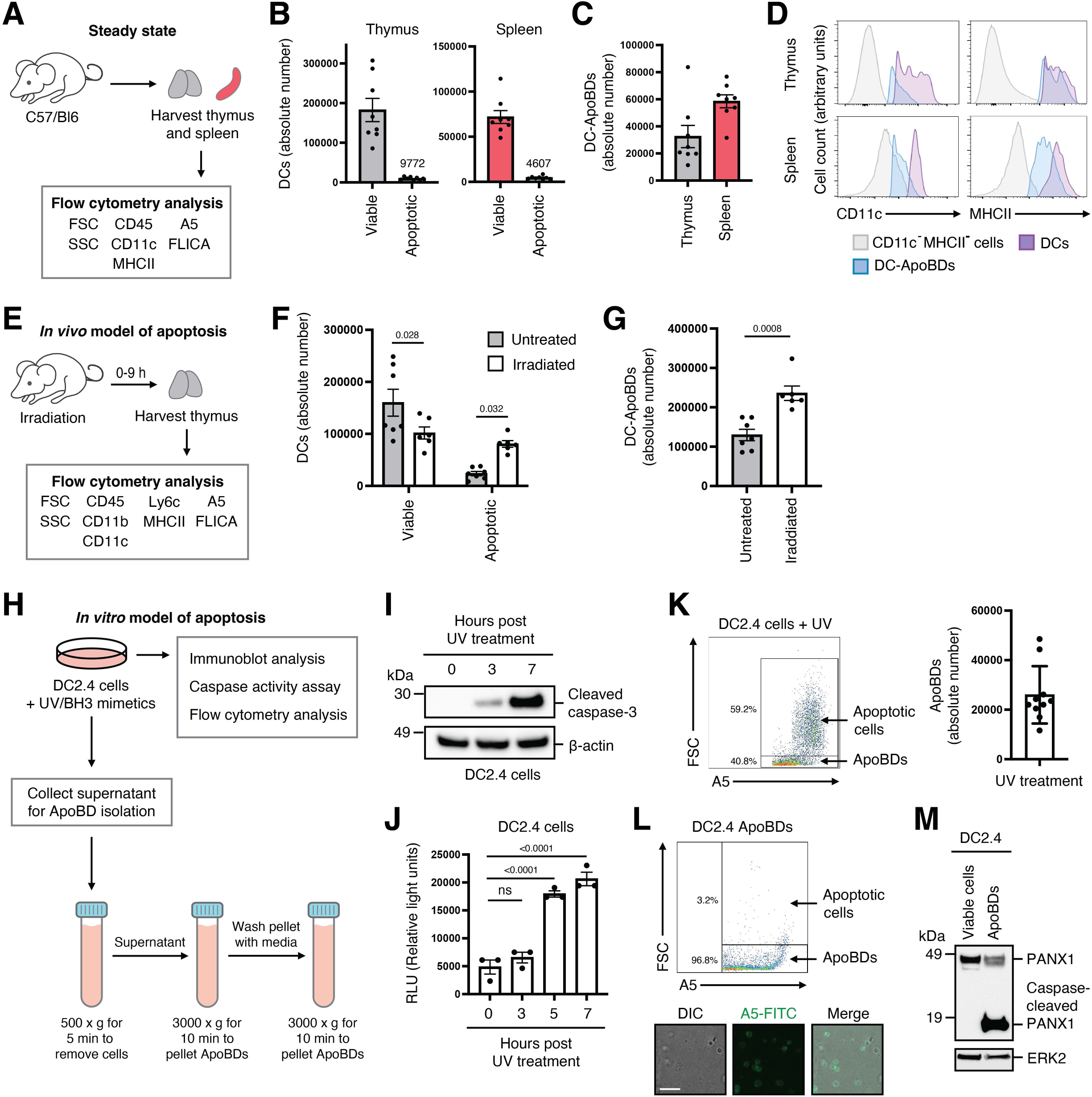
Generation of DC-derived ApoBDs under *in vivo* and *in vitro* conditions. **(A)** Schematic diagram to determine the levels DC-derived ApoBDs in the spleen and thymus of 10 weeks old C57/Bl6 under homeostatic conditions (see Supplementary Figure 1A and B for electronic gating strategy). **(B)** Absolute numbers of viable and apoptotic (A5^+^ FLICA^+^) DCs in the thymus and spleen (*n*= 8). **(C)** Absolute number of DC-ApoBDs in the thymus and spleen (*n*= 8). **(D)** Flow cytometry analysis of the levels of CD11c and MHCII surface markers on DC-ApoBDs (blue) and viable DCs (purple), with CD11c^−^ MHCII^−^ immune cells (grey) as a comparison. **(E)** Schematic diagram of the *in vivo* mouse model of whole-body X-ray irradiation-induced apoptosis (680 rad, see Supplementary Figure 1A and B for electronic gating strategy). Absolute number of viable and apoptotic DCs **(F)** and DC-ApoBDs **(G)** in the thymus of untreated (*n*=7) and 9 h post-irradiation treated mice (*n*=6), was determined via flow cytometry. **(H)** Schematic diagram of the *in vitro* model of apoptosis utilising the DC2.4 cell line and the isolation of DC-ApoBDs from culture supernatant by differential centrifugation. **(I)** Immunoblot analysis of caspase 3 activation in UV irradiated (150 mJ/cm^2^) DC2.4 cells (*n*=3). **(J)** Caspase 3/7 activation in DC2.4 cells 0-6 h post UV irradiation, as determined by caspase 3/7 activity assay (*n*=3). **(K)** Flow cytometry analysis of DC2.4 apoptotic cells and ApoBDs following UV irradiation. 1.5×10^5^ DC2.4 cells were induced to undergo apoptosis (*n*=10). **(L)** Purity of DC2.4 ApoBDs isolated via differential centrifugation was determined by flow cytometry and confocal microscopy analysis (*n*=3). **(M)** Immunoblot analysis of full length PANX1 and caspase-cleaved PANX1 in viable DC2.4 cells and ApoBDs isolated from apoptotic DC2.4 cells. Error bars represent s.e.m. (*n*= individual mice or independent experiments). Statistical analysis for (B, C, F, G) was performed using unpaired Student’s two-tailed *t*-test, ns= P>0.05. Statistical analysis for (J) was performed using two-way ANOVA with Sidak’s multiple comparisons test, ns= P>0.05.

To determine whether we could recapitulate the generation of DC-derived ApoBDs from apoptotic DCs *in vitro*, a DC-like cell line DC2.4 was used. First, DC2.4 cells were induced to undergo apoptosis via ultraviolet (UV) irradiation or treatment with BH3 mimetics (ABT-737 and S63845, inhibitors of pro-survival BCL-2 and MCL-1, respectively) (Figure 1H). Induction of DC2.4 cell undergoing apoptosis was confirmed by caspase 3 activation as determined by immunoblot analysis and caspase 3/7 activity assay (Figure 1I and J). Formation of DC-ApoBDs in the cultured supernatant following UV treatment was also confirmed by flow cytometry analysis (Figure 1K, Supplementary Figure 2A). To characterise ApoBDs generated by DCs based on our previously established criteria for apoptotic EV characterisation (Poon *et al*, 2019), we isolated DC-ApoBDs from apoptotic DC2.4 cells by differential centrifugation (Figure 1H). Purity of isolated DC-ApoBDs and the exposure of the apoptotic marker PtdSer was confirmed by flow cytometry and confocal microscopy analysis (Figure 1L). DC-ApoBDs were ∼2.5 μm in diameter (Supplementary Figure 2B). The presence of caspase-cleaved proteins such as PANX1 in DC-ApoBDs was also validated by immunoblot analysis (Figure 1M). Collectively, these data suggest apoptotic DCs can generated ApoBDs in *in vivo* and *in vitro* settings.

### Apoptotic DCs undergo disassembly via distinct morphological steps

The formation of ApoBDs is regulated by distinct morphological steps, namely membrane blebbing, the generation of a thin string-like membrane protrusion called apoptopodia, and finally fragmentation into distinct ApoBDs. Notably, different cell types can undergo apoptotic cell disassembly via different morphological steps (Atkin-Smith & Poon, 2017). Thus, to characterise the morphological steps involved in the disassembly of apoptotic DCs, time-lapse confocal microscopy was performed. The majority of DC2.4 cells induced to undergo apoptosis by UV irradiation first underwent surface and dynamic membrane blebbing, and subsequently the formation of apoptopodia during the progression of apoptosis (Figure 2A and B). Morphological changes of DC2.4 cells undergoing apoptosis was also captured by transmission electron microscopy (Figure 2C). Exposure of the ‘eat-me’ signal and apoptosis marker PtdSer during the disassembly of apoptotic DC2.4 cells was confirmed by confocal microscopy (Figure 2D). It should be noted that the morphological hallmarks of apoptotic cell disassembly as described above were also observed when DC2.4 cells were induced to undergo apoptosis by BH3 mimetics (Supplementary Figure 3A), as well as primary human monocyte-derived DCs (HMDCs) induced to undergo apoptosis by UV irradiation (Figure 2E). To examine whether activated DCs can also undergo similar morphological changes to generate ApoBDs, DC2.4 cells were activated by LPS as confirmed by the upregulation of activation marker CD40 and CD80 (Figure 2F), and similar apoptotic morphologies were observed (Figure 2G). Together, these data indicate DCs can undergo distinct morphological steps during apoptosis to generate ApoBDs.

**Figure 2.**
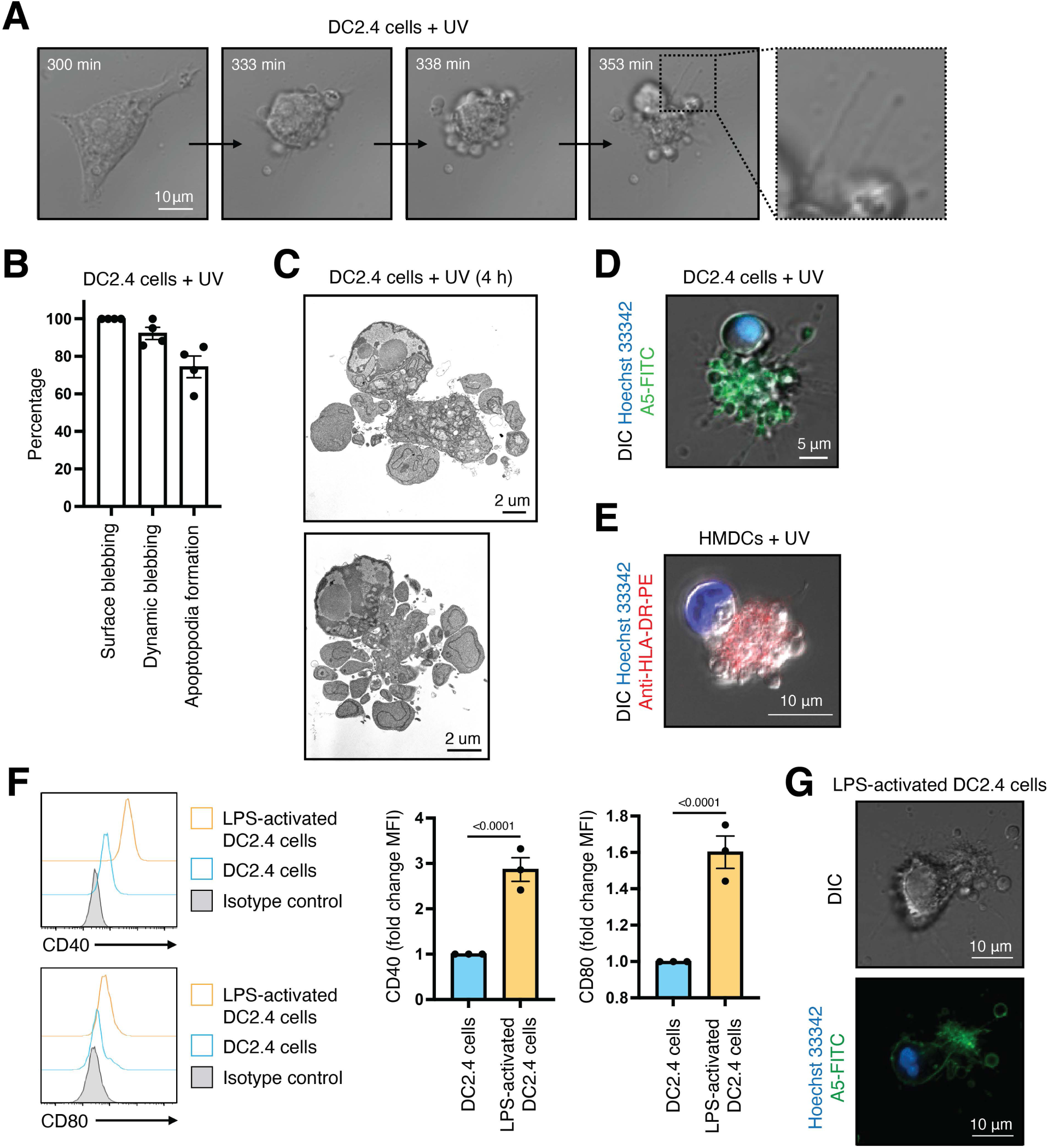
Apoptotic DCs generate ApoBDs via membrane blebbing and apoptopodia formation. **(A)** Time-lapse DIC microscopy images monitoring the formation of membrane blebs and apoptopodia by DC2.4 cells treated with UV (150 mJ/cm^2^) to induce apoptosis. **(B)** Quantitation of live microscopy data from (A) to determine the percentage of apoptotic DC2.4 cells that underwent surface blebbing, dynamic blebbing, and apoptopodia formation (*n*=4). **(C)** Transmission electron microscopy images of apoptotic DC2.4 cells treated with UV to induce apoptosis. **(D)** A5-FITC and Hoechst 33342 staining of PtdSer and nuclear DNA, respectively, on/in DC2.4 cell induced to undergo apoptosis by UV irradiation. **(E)** Anti-HLA-DR-PE and Hoechst 33342 staining of MHCII and nuclear DNA, respectively, on/in HMDCs induced to undergo apoptosis by UV irradiation. **(F)** Flow cytometry analysis of the level of CD40 and CD80 on DC2.4 cells and LPS-activated DC2.4 cells. **(G)** A5-FITC and Hoechst 33342 staining of LPS-activated DC2.4 cell induced to undergo apoptosis by UV irradiation. Error bars represent s.e.m. Data are representative of at least three independent experiments. Statistical analysis for (F) was performed using unpaired Student’s two-tailed *t*-test,

### Formation of DC-ApoBDs is regulated by ROCK1 kinase and T-type calcium channels

A number of molecular factors including ROCK1, pannexin 1 (PANX1) and T-type calcium channels have been described to regulate the formation of ApoBDs in cell types such as T cells, monocytes, fibroblasts, and epithelial cells (Tixeira *et al*, 2020; Poon *et al*, 2014; Atkin-Smith *et al*, 2015; Phan *et al*, 2023). Thus, the importance of these apoptotic cell disassembly machineries in regulating DC-ApoBD formation were investigated further. ROCK1 kinase is activated by caspase-mediated cleavage and drives the formation of membrane blebs during apoptosis by promoting actin-myosin contraction (Coleman *et al*, 2001; Sebbagh *et al*, 2001). We first confirmed the presence of caspase-activated fragment of ROCK1 in DC2.4 cells induced to undergo apoptosis by UV irradiation (Figure 3A). Similar to previous studies (Tixeira *et al*, 2020), inhibition of ROCK1 using the ROCK inhibitor GSK-269962 blocked the formation of dynamic membrane blebs but not surface blebs by apoptotic DC2.4 cells (Figure 3B-D). Furthermore, GSK-269962 treatment also inhibited the formation of DC-ApoBDs (Figure 3E) without affecting the overall level of apoptosis (Figure 3F) or the exposure of PtdSer on ApoBDs and apoptotic cells (Figure 3G). Together, these studies demonstrate ROCK1 is a positive regulator of DC-ApoBD formation.

**Figure 3.**
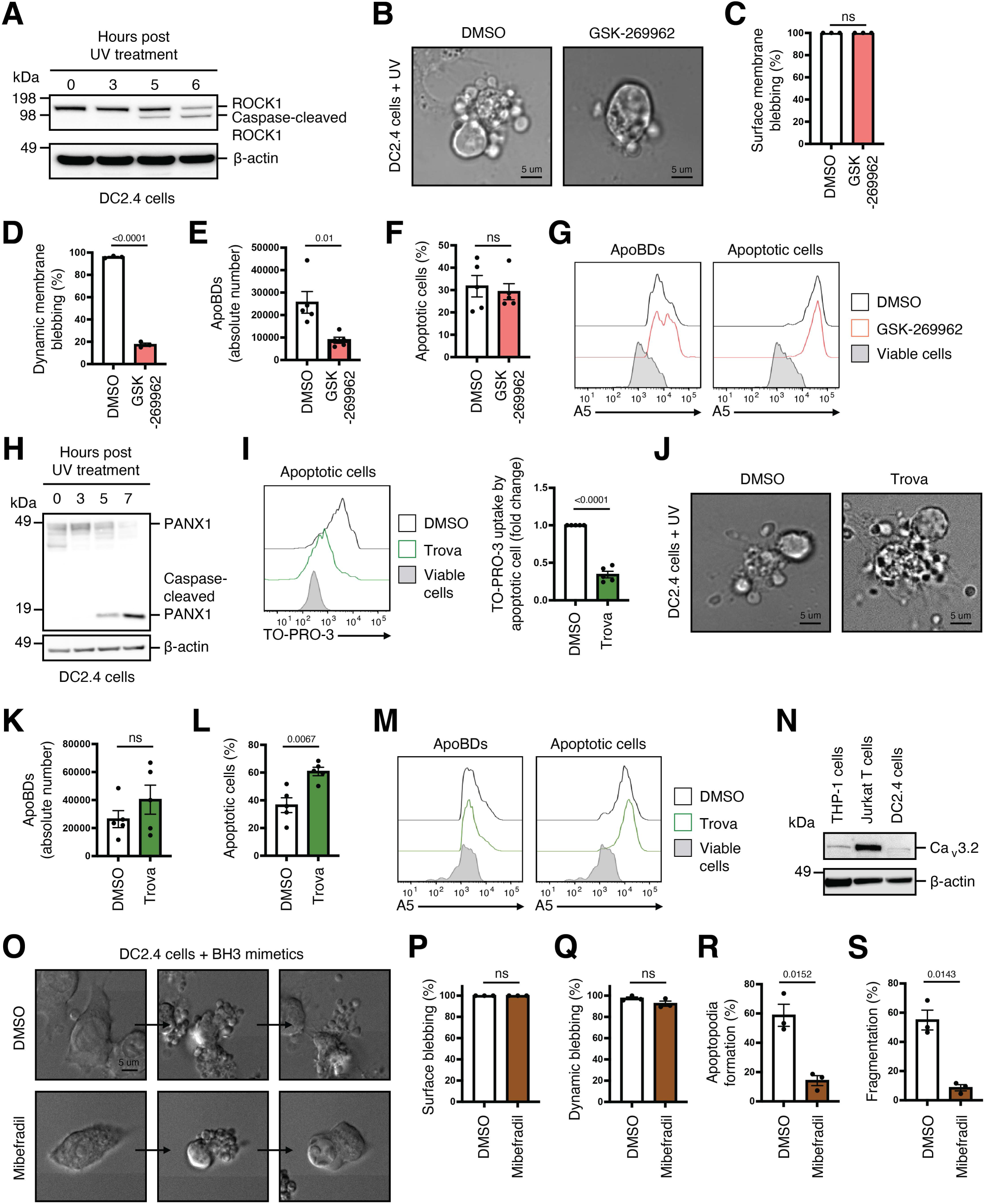
DC disassembly during apoptosis is regulated by ROCK1 and T-type calcium channels, but not PANX1. **(A)** ROCK1 protein expression and proteolytic processing in untreated and apoptotic samples (UV-treated, 150 mJ/cm^2^) was determined by immunoblotting. **(B)** DIC microscopy images monitoring membrane blebbing of DC2.4 cells treated with UV to induce apoptosis and in the presence of vehicle control (DMSO) or the ROCK1 inhibitor GSK-269962 (1 μM). Quantitation of live microscopy data from (B) to determine the percentage of apoptotic DC2.4 cells that underwent surface blebbing **(C)** and dynamic blebbing **(D)** (*n*=3). Formation of ApoBDs **(E)** and apoptotic cells **(F)** by DC2.4 cells treated with UV to induce apoptosis and in the presence of DMSO or GSK-269962 (1 μM). 1.5×10^5^ DC2.4 cells were induced to undergo apoptosis (*n*=5). **(G)** Flow cytometry analysis PtdSer exposure on ApoBDs and apoptotic cells generated from UV-treated DC2.4 cells in the presence of DMSO or GSK-269962 (1 μM), based on A5-FITC staining. **(H)** PANX1 protein expression and proteolytic processing in untreated and apoptotic samples (UV-treated) was determined by immunoblotting. **(I)** Flow cytometry analysis of TO-PRO-3 dye uptake by apoptotic cells generated from UV-treated DC2.4 cells in the presence of vehicle control (DMSO) or the PANX1 inhibitor trovafloxacin (Trova, 20 μM) (*n*=5). **(J)** DIC microscopy images monitoring apoptopodia formation by DC2.4 cells treated with UV to induce apoptosis and in the presence of DMSO or Trova (20 μM). Formation of ApoBDs **(K)** and apoptotic cells **(L)** by DC2.4 cells treated with UV to induce apoptosis and in the presence of DMSO or Trova (20 μM). 1.5×10^5^ DC2.4 cells were induced to undergo apoptosis (*n*=5). **(M)** Flow cytometry analysis of PtdSer on ApoBDs and apoptotic cells generated from UV-treated DC2.4 cells in the presence of DMSO or Trova (20 μM), based on A5-FITC staining. **(N)** Ca_v_3.2 T-type calcium protein expression in human THP-1, Jurkat T cells and DC2.4 cells was determined by immunoblotting. **(O)** Time-lapse DIC microscopy images monitoring the formation of membrane blebs and apoptopodia by DC2.4 cells treated with BH3 mimetics (ABT-737, 2 μM; S63845, 5 μM) to induce apoptosis and in the presence of vehicle control (DMSO) or the T-type calcium channel inhibitor mibefradil (10 μM). Quantitation of live microscopy data from (O) to determine the percentage of apoptotic DC2.4 cells that underwent surface blebbing **(P)**, dynamic blebbing **(Q)**, apoptopodia formation **(R)**, and cell fragmentation **(S)** (*n*=3). Error bars represent s.e.m. Statistical analysis was performed using unpaired Student’s two-tailed *t*-test, ns= P>0.05. Data are representative of at least three independent experiments.

Next, we investigated the role of caspase-activated PANX1 membrane channels, a key negative regulator of apoptopodia and ApoBD formation for T cells (Poon *et al*, 2014), in the disassembly of apoptotic DCs. Caspase-mediated processing and activation of PANX1 in apoptotic DC2.4 cells was confirmed by immunoblot analysis (Figure 3H) and the uptake of the DNA-binding dye TO-PRO-3, which enters early apoptotic cells via PANX1 (Chekeni *et al*, 2010), into apoptotic DC2.4 cells was determined by flow cytometry (Figure 3I). Furthermore, uptake of TO-PRO-3 into apoptotic DC2.4 cells was inhibited by the PANX1 inhibitor trovafloxacin (trova) (Figure 3I), indicating PANX1 is activated in apoptotic DCs. Notably, the level of PANX1 activation is relatively lower for apoptotic DC2.4 cells compared to apoptotic Jurkat T cells (Supplementary Figure 3B). Since PANX1 is a negative regulator of apoptopodia and ApoBD formation, blockade of PANX1 by trova has been shown to promote apoptopodia and ApoBD formation in cell types including T cells (Poon *et al*, 2014). Although PANX1 is activated in apoptotic DC2.4 cells (Figure 3H and I), trova treatment did not have an obvious impact on the formation of apoptopodia and ApoBDs by DC2.4 cells undergoing apoptosis (Figure 3J-M), possibly reflecting the relatively lower level of PANX1 activation in apoptotic DC2.4 cells and the ability of DC2.4 cells to form an abundance of apoptopodia and ApoBDs under normal conditions (Figure 1K). Collectively, PANX1 is not an important regulator of apoptotic DCs disassembly.

Recently, we have identified T-type calcium channels such as Ca_v_3.2 as a positive regulator of apoptopodia and ApoBD formation, whereby inhibition of T-type calcium channels pharmacologically with mibefradil blocks apoptotic cell disassembly of cell types including Jurkat T cells, THP-1 cells and Vero E6 epithelial cells (Phan *et al*, 2023). Expression of Ca_v_3.2 in DC2.4 cells was confirmed by RT-qPCR and immunoblot analysis (Supplementary Figure 2C, Figure 3N). Importantly, mibefradil treatment markedly reduced the formation of apoptopodia and fragmentation of apoptotic DC2.4 cells without affecting the membrane blebbing stages of apoptotic cell disassembly as determined by time-lapse microscopy (Figure 3O-S). Collectively, these data demonstrate the formation of DC-ApoBDs by apoptotic DC2.4 cells is regulated by ROCK1 and T-type calcium channels but not PANX1. To further evaluate the importance of ROCK1 and T-type calcium channels in the formation of ApoBDs by primary DCs, BMDCs were treated with GSK-269962 or mibefradil, respectively. BMDCs induced to undergo apoptosis by X treatment exhibited similar morphological changes as apoptotic DC2.4 cells, in particular the formation of numerous apoptopodia dynamically during apoptosis progression (Supplementary Figure 4A). Under these apoptotic conditions, treatment with mibefradil but not GSK-269962 markedly inhibited the fragmentation of BMDCs (Supplementary Figure 4B), suggesting T-type calcium channels but not ROCK1 is important for apoptotic BMDC disassembly, possibly due to the propensity for BMDCs to generate a large amount of apoptopodia.

### DC-ApoBDs aid direct antigen presentation

DCs play a key role in regulating adaptive immunity by mediating antigen presentation to different T cell subsets. To facilitate direct antigen presentation, DCs display antigen presentation machineries including MHCI or MHCII and co-stimulatory molecules on the cell surface (Hilligan & Ronchese, 2020; Théry & Amigorena, 2001). To examine the possibility that DC-ApoBDs can elicit T cell activation and proliferation through direct antigen presentation, we first confirmed the presence of MHCI and co-stimulatory molecules such as CD80 and CD86 on DC-ApoBDs generated from LPS-activated apoptotic DC2.4 cells (Figure 4A). Next, to determine if DC-ApoBDs can present antigen to naïve T cells, we utilised the OT.I transgenic system whereby CD8^+^ T cells from OT.I transgenic mice were restricted to the MHCI subtype H2K^b^ and SIINFEKL_257-264_ (OVA antigenic peptide) (Clarke *et al*, 2000). CD8^+^ OT.I T cells were subsequently co-cultured with ApoBDs derived from BMDCs expressing H2K^b^ pulsed with the SIINFEKL peptide (Figure 4B), as characterised below. We first confirmed the ability of apoptotic BMDCs to generate ApoBDs following apoptosis induction with UV irradiation and pulsing BMDCs with SIINFEKL or a non-OT.I specific peptide (gBT-1_498-505_) prior to induction of apoptosis did not affect the level of ApoBD formation by BMDCs (Supplementary Figure 5A).

**Figure 4.**
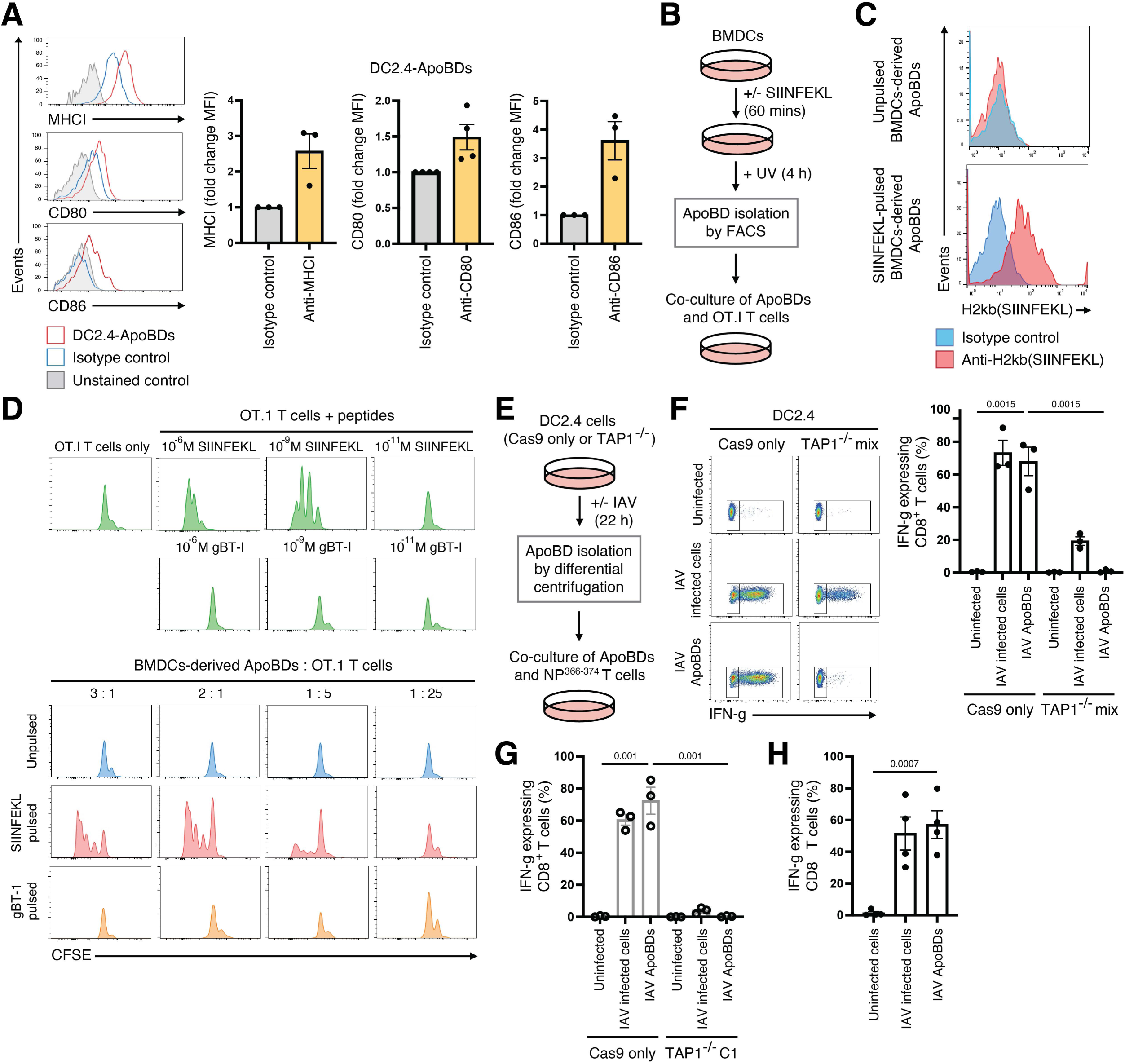
DC-ApoBDs contain biomolecules for antigen presentation. **(A)** Flow cytometry analysis of the level of MHCI, CD80 and CD86 on ApoBDs generated from LPS-activated DC2.4 cells, compared to isotype control (mouse IgG2a kappa, hamster IgG2 kappa and rat IgG2a kappa, respectively) and unstained control. Apoptosis was induced by UV treatment (150 mJ/cm^2^) (*n*=3-4). **(B)** Schematic of the co-culture assay of CD8^+^ T cells obtained from OT.I transgenic mice with FACS-isolated BMDC-derived ApoBDs. **(C)** The level of H2Kb(SIINFEKL) complex on ApoBDs generated from unpulsed or SIINFEKL-pulsed BMDCs was measured by flow cytometry, with IgG1 as an isotype control. **(D)** Representative histograms of CFSE-stained naïve OT.I T cells cultured in the presence of different concentrations of SIINFEKL or gBT-1 peptides, or the indicated ratio of ApoBDs derived from unpulsed, SIINFEKL-pulsed or gBT-I-pulsed BMDCs (*n*=2). **(E)** Schematic of the co-culture assay of NP^366-374^ specific CD8^+^ T cells with ApoBDs isolated by differential centrifugation from IAV-infected Cas9 only or TAP1^−/−^ DC2.4 cells. IFNγ expression by NP^366-374^ specific CD8^+^ T cells cocultured with uninfected cells, IAV-infected cells and ApoBDs isolated from IAV-infected Cas9 only or TAP1^−/−^ mix DC2.4 cells **(F)** or TAP1^−/−^ clone 1 (C1) DC2.4 cells **(G)** was quantified by flow cytometry (*n*=3). **(H)** IFNγ expression by NP^366-374^ specific CD8^+^ T cells cocultured with uninfected cells, IAV-infected cells and FACS-isolated ApoBDs from IAV-infected Cas9 only DC2.4 cells (*n*=4). Error bars represent s.e.m. Statistical analysis was performed using unpaired Student’s two-tailed *t*-test. Data are representative of at least three independent experiments unless otherwise stated.

Furthermore, to ensure that MHCI in complex with the antigenic SIINFEKL peptide was present on the surface of BMDC-derived ApoBDs after peptide pulsing, an antibody against the H2K^b^(SIINFEKL_257-264_) complex was used for flow cytometry analysis and confirmed MHCI/peptide complex was detectable on the surface of SIINFEKL-pulsed BMDC-derived ApoBDs (Figure 4C). Next, BMDC-derived ApoBDs isolated by fluorescence activated cell sorting (Supplementary Figure 5B) were co-cultured with CFSE-stained CD8^+^ OT.I T cells for 72 hours (Figure 4B), with a reduction in CFSE fluorescence indicative of T cell proliferation (Quah *et al*, 2007). As a control, CFSE-stained naïve CD8^+^ OT.I T cells cultured in the presence of increasing concentrations of the SIINFEKL peptide (10^−11^ M to 10^−6^ M) but not the non-OT.I specific peptide gBT-1_498-505_ showed proliferation proportional to the concentration of SIINFEKL peptide in culture (Figure 4D). Importantly, SIINFEKL-pulsed BMDC-derived ApoBDs induced T cell proliferation, but not ApoBDs derived from unpulsed or gBT-1-pulsed BMDC-derived ApoBDs (Figure 4D). The level of T cell proliferation correlated with the number of ApoBDs in the co-culture, with the highest ratio of SIINFEKL-pulsed BMDC-derived ApoBDs to naïve CD8^+^ OT.I T cells resulted in the highest level of T cell proliferation (Figure 4D). Collectively, these data demonstrate the ability of DC-ApoBDs to present antigen directly and prime naïve T cells.

To further examine the ability of DC-ApoBDs in mediating direct antigen presentation through MHCI-peptide complexes, we generated TAP1 deficient DC2.4 cells via CRISPR-Cas9 system (Supplementary Figure 6A), which would block the trafficking of cytoplasmic peptides into the ER lumen for MHCI loading (Hewitt, 2003), thus impairing the presentation of antigens via MHCI. Next, we utilised influenza A virus (IAV) infection model to test the ability of ApoBDs generated from IAV-infected apoptotic DC2.4 cells to activate IAV-specific CD8^+^ T cells (Figure 4E). We first confirmed that both the Cas9 control and TAP1 deficient DC2.4 cells undergo apoptosis and apoptotic cell disassembly to generate DC-ApoBDs following IAV infection (Supplementary Figure 6B-D). To monitor the ability of DC-ApoBDs to activate T cells, we quantified the level of IFN-γ producing CD8^+^ T cells (NP_366-374_) following co-culture with IAV infected Cas9 or TAP1^−/−^ mix DC2.4 cells or ApoBDs isolated from these cells (Figure 4F). CD8^+^ T cells co-cultured with DC-ApoBDs generated from Cas9 but not TAP1^−/−^ mix DC2.4 cells lead to IFN-γ production, to a level comparable to CD8^+^ T cells co-cultured with IAV-infected DC2.4 cells (Figure 4F). As we noted a small proportion of IFN-γ producing T cells when co-cultured with IAV-infected TAP1^−/−^ mix DC2.4 cells, possibly due to the presence of TAP1 sufficient cells in the mixed population, we also generated clonal TAP1^−/−^ DC2.4 cells (Supplementary Figure 6A) and showed that DC-ApoBDs generated from Cas9 but not TAP1^−/−^ C1 DC2.4 cells lead to IFN-γ production (Figure 4G). To further ascertain that DC-ApoBDs generated from IAV-infected DC2.4 cells can activate cognate CD8^+^ T cells, DC-ApoBDs were isolated by FACS to high purity and found to promote IFN-γ production (Supplementary Figure 6E, Figure 4H), to a comparable level as DC-ApoBDs isolated through differential centrifugation (Figure 4G). Altogether, these data demonstrate the ability of DC-ApoBDs to present antigen directly to T cells via a MHC-restricted TAP1-dependent mechanism in a viral infection setting.

## Discussion

Small EVs such as exosomes (30-100 nm in diameter) and microvesicles (50-1,000 nm) have been shown to exhibit immune modulation properties via a variety of mechanisms (Buzas, 2023), including exosomes transferring miRNA at the immunological synapse (Mittelbrunn *et al*, 2011), the transfer of key pattern recognition molecules such as toll-like receptors (TLRs) between cells via BMDC-derived exosomes (Zhang *et al*, 2019), and microvesicles stimulating IgG production by B cells (J & Cm, 2006). Amongst the many functions of EVs in modulating immune responses, EVs have also been shown to play a role in antigen presentation pathways whereby key antigen presentation molecules such as MHCI and MHCII can be found as MHC-peptide complexes on exosomes, and these EVs can mediate priming of CD4^+^ and CD8^+^ T cells leading to their activation (Tkach *et al*, 2017; Théry *et al*, 2002; Segura *et al*, 2005; Utsugi-Kobukai *et al*, 2003; Admyre *et al*, 2006). Furthermore, MHCII can be found in small EVs isolated from spleen-derived murine DC line (D1) induced to undergo apoptosis by UV irradiation (Théry *et al*, 2001). There is also evidence of exosomes and apoptotic cell-derived EVs facilitating the transfer of antigenic peptide to DCs as a means of antigen presentation (Schnitzer *et al*, 2010; Schaible *et al*, 2003). Our finding that ApoBDs, a subtype of large EVs derived from apoptotic cells, can also mediate direct antigen presentation via functional MHC-peptide complexes demonstrates for the first time that following cell death, EVs generated from antigen presenting cells such as DCs can continue to directly regulate antigen-specific MHC-restricted T cell responses, resulting in T cell proliferation and activation. Although it is well described that apoptotic cells can carry antigens and the uptake of apoptotic materials by DCs can subsequently lead to cross-presentation (Blachère *et al*, 2005; Albert *et al*, 1998), the ability of DC-derived ApoBDs to directly present antigens to naïve and pre-activated antigen-specific T cells raises a number of interesting possibilities. First, the removal of DCs after antigen presentation by T cells via a perforin and granzyme-dependent mechanism is well documented and is generated in response to virus-mediated DC killing in peripheral tissues or infected lymph nodes as a part of an essential mechanism for T cell regulation. Classically, the removal of DCs after antigen presentation is seen as a mechanism of negative feedback regulation in which the killing of DCs occurs to remove the initiating stimulus and prevent interminable expansion of antigen-specific T cells (Yang *et al*, 2006; Belz *et al*, 2007). The ability of DC-ApoBDs to continue to engage with T cells and directly present antigen resulting in T cell activation post apoptosis induction suggests that such feedback regulation may occur only when all DC-derived apoptotic materials are removed by phagocytes. Furthermore, DC-ApoBDs bearing MHC-peptide complexes may also facilitate antigen presentation to help sustain an adaptive immune response through the cross-presentation pathway or via cross-dressing. Notably, the formation of ApoBDs has been shown recently to aid the uptake of apoptotic materials by DCs (Tixeira *et al*, 2020) and CD8^+^ DCs are excellent at ingesting apoptotic materials and cross-presenting this material via MHCI (Janssen & Thacker, 2012). Thus, it is conceivable that ApoBDs could also facilitate antigen presentation through a recipient DC reprocessing antigens derived from DC-ApoBDs or utilising MHC-peptide complexes and co-stimulatory molecules on DC-ApoBDs through the cross-presentation and cross-dressing pathways, respectively. Furthermore, our findings show that DC-derived ApoBDs are formed in the steady state. Notably, steady state maintenance of DC levels via apoptosis is imperative to avoid downstream immune dysregulation, with previous studies demonstrating that overexpression of caspase inhibitors or preventing Fas-mediated death in DCs leads to DC accumulation and the development of systemic autoimmunity (Chen *et al*, 2006, 2007). Thus, DC apoptosis is important for maintaining self-tolerance and whether DC-ApoBDs generated during steady state could also participate in this immune regulation process warrants further investigation.

It is becoming more apparent that different cell types can generate ApoBDs via distinct mechanisms and can regulate different processes based on the contents these ApoBDs carry. For example, apoptotic monocytes generate ApoBDs via the fragmentation of a distinct membrane protrusion known as beaded-apoptopodia (Atkin-Smith *et al*, 2015). Furthermore, influenza A virus (IAV) infected monocytes undergoing apoptosis can form ApoBDs that carry viral particles and the trafficking of these ApoBDs to neighbouring cells propagates IAV infection (Atkin-Smith *et al*, 2020). Disassembly of apoptotic endothelial cells under inflammatory conditions can also aid the transfer of inflammatory cytokine IL-α via ApoBDs (Berda-Haddad *et al*, 2011). Similarly, the formation of ApoBDs by apoptotic mature osteoclasts can mediate the trafficking of cell surface molecules like RANKL to regulate intercellular communication (Ma *et al*, 2019). Notably, how endothelial cells and osteoclasts undergo fragmentation during apoptosis has not been characterised in detail. In this study, we described the ability of DCs to generate ApoBDs via the formation of dynamic membrane blebs and apoptopodia. Moreover, ROCK1 for DC2.4 cells and T-type calcium channels for DC2.4 and BMDCs were identified as molecular regulators of DC-ApoBD formation by controlling the aforementioned morphological processes, respectively. It should be noted that similar to apoptotic monocytes but in contrast to apoptotic T cells, DCs undergoing apoptosis readily form apoptopodia and ApoBDs under basal settings *in vitro*, which correlated inversely with the levels of PANX1 activation (Supplementary Figure 3B). These observations further highlight that the activities of apoptotic cell disassembly regulators could vary between different cell types, resulting in the differential dependency on membrane blebbing and apoptopodia to generate ApoBDs. Moreover, it would of be interest in future studies to examine whether modulation of these regulatory factors specifically in DCs could influence immune responses, in particular direct antigen presentation in steady state and in infection settings.

Collectively, this work has generated novel insights into the formation of large EVs by apoptotic DCs as well as the underpinning mechanisms. The described role of DC-ApoBDs in triggering specific CD8^+^ T cell response through direct antigen presentation further highlights the importance of ApoBD formation in mediating immune cell communication following cell death.

## Materials and Methods

### Reagents

RPMI-1640 medium was purchased from Life Technologies. Foetal bovine serum (FBS), Hoechst 33342, TO-PRO-3 iodide, penicillin and streptomycin from Thermofisher Scientific. MycoZap reagent was purchased from Lonza, and EDTA from Sigma-Aldrich. The following reagents were purchased from BD Biosciences: rat anti-mouse CD16/32, anti-mouse CD45.2-BV421, anti-mouse Ly6C-BV421, anti-mouse CD45.2 A700, anti-mouse CD11c-PE, anti-mouse IFN-γ-FITC, anti-mouse CD8-APC, anti-mouse CD3-PECy7, anti-mouse B220-PECy7, anti-mouse CD11b-PECy7, anti-mouse A-I/E-I-PerCp5.5, anti-mouse A-I/E-I-BV650, A5 binding buffer, A5-APC, A5-BV421, A5-V450 and A5-FITC.

### Mice

Female and male C57Bl/6 mice were used for all *in vivo* models. 4–6-week-old mice were used for the X-ray irradiation model and 10-12-week-old mice were used for steady state analysis. All mice were used under approval of AEC21034, La Trobe University. All experiments were approved by the La Trobe University Animal Ethics Committee and in accordance with the National Health and Medical Research Council Australia code of practice for the care and use of animals for scientific purposes. Sample size was determined based on previous studies (Poon *et al*, 2014) and no blinding was performed.

### Analysis of the spleen and thymus in steady state

To determine homeostatic levels of apoptosis, spleens and thymi were harvested from untreated 10-12-week-old mice. The thymus was harvested and mechanically cut with scissors then digested enzymatically with collagenase type I (Thermofisher Scientific) diluted in complete RPMI-1640 medium for 20 min at room temperature with gentle resuspension. Spleens were dissociated mechanically and splenocytes were treated with Red Blood Cell lysis buffer twice. All samples were filtering through a 70 μm cell strainer, resuspended in complete RPMI and centrifuged at 3,000 x *g* for 6 min to sediment cells and ApoBDs. Samples were stained with Vybrant FAM Caspase 3 and 7 assay kit with a fluorescent inhibitor of caspases (FLICA) (Invitrogen) following the manufacturer’s protocol. Samples were treated with rat-anti-mouse CD16/32 (1:300, clone 2.4G2) at 4°C for 10 min followed by staining with A5-APC (1:200) in 1x A5 binding buffer for 10 min at room temperature. Samples were then stained with immune cell antibody panel consisting of CD45.2-BV421, CD11c-PE, CD3-PECy7, B220-PECy7, A-I/E-I-PerCp5.5 and incubated at 4°C for 20 min. All samples were analysed by flow cytometry using the BD FACS CANTO II (BD Biosciences) and absolute number of cells and ApoBDs were determined using Sphero AccuCount Particles (ProSciTech).

### Irradiation mouse model

To induce apoptosis *in vivo*, 4–6-week-old mice were treated with a dose of 680 rad of X-ray irradiation using the Rad Source RS-2000 Irradiator (Rad Source Technologies Inc). Mice were euthanised by CO_2_ asphyxiation 9 hours post-treatment. The thymus and single cell suspension was prepared as per above. Samples were treated with rat-anti-mouse CD16/32 (1:300) followed by staining with A5-BV605 in 1x A5 binding buffer as described previously. Samples were then stained with immune cell antibody panel consisting of CD45.2-A700, CD11c-PE, CD11b-PECy7, Ly6C-BV421, A-I/E-I-BV650 and incubated at 4°C for 20 min. All samples were analysed by flow cytometry using the Cytoflex S (Beckman Coulter) and absolute number of cells and ApoBDs was determined using Sphero AccuCount Particles (ProSciTech).

### Generation of HMDCs and BMDCs

To generated HMDCs, fresh buffy coat was obtained the Australia Red Cross Blood Service (Melbourne, Australia) with an appropriate ethics approval (FHEC09/16) from La Trobe Human Ethics Committee. Peripheral blood mononuclear cells were collected by Ficoll gradient and CD14^+^ monocytes were obtained by positive magnetic isolation using a MACS LS column with a MACS separator (Miltenyi Biotech) in accordance with the manufacturer’s instructions. CD14^+^ monocytes adhered to culture flask and cultured in complete RPMI-1640 containing 10% FCS, 1,000 U/mL human GM-CSF (Miltenyi Biotech), and 400 U/mL human IL-4 (Miltenyi Biotech) for the generation of DCs. HMDCs (cells in suspension) were used on the seventh day. To generated BMDCs, bone marrow cells was harvested from C57Bl/6 mice and cultured in RPMI-1640/10%FSC containing 10% of X63-GM-CSF supernatant (containing 10 ng/mL GM-CSF) as per our previous study (Deng *et al*, 2021). Fresh culture medium was added on day 3 and 6, and BMDCs (cells in suspension) were used on the eighth day.

### Cell culture

DC2.4 cells, THP-1 and Jurkat T cells (ATCC) were cultured in complete RPMI media, constituted of RPMI-1640 supplemented with 10% FBS, 50 IU/mL penicillin and 50 µg/mL streptomycin. All cells were maintained in a humidified environment at 37°C with 5% CO_2_. Cells are routinely tested for mycoplasma contamination.

### CRISPR/Cas9 gene editing

CRISPR/Cas9 technology was used as per previous studies (Tixeira *et al*, 2020) to generate a genetic disruption of the TAP1 gene in DC2.4 cells. The following oligonucleotides correspond to gRNA sequences targeting the TAP1 gene: 5’TCCCAATGGCCATTCCCTTCTTCA-3’, 5’AAACTGAAGAAGGGAATGCCATT-3’, and were designed through the website https://www.synthego.com/products/bioinformatics/crispr-design-tool. To induce gRNA synthesis, cells were treated with doxycycline (1 µg/mL) and bulk sorted by FACS based on the expression of mCherry (indicative of Cas9 expression) and GFP (indicative of gRNA expression). Single-cell sorting was then done manually into individual wells of a 96 well plate and cells were sorted again based on mCherry and GFP expression to collect a TAP1 knockout clone.

### Induction of apoptosis *in vitro*

Cells were induced to undergo apoptosis in serum-free RPMI media supplemented with 1% BSA (SF-RPMI/BSA). Cells were treated with either BH3 mimetics ABT-737 (2 μM) and S63845 (5 μM) (MedChemExpress) or UV irradiation (150 mJ/cm^2^) using the Stratagene Stratalinker 1800 (Agilent Technologies) to induce apoptosis. Following apoptosis induction, cells were incubated for the indicated duration at 37°C with 5% CO_2_ before further use or analysis.

### Caspase 3/7 activity assay

Caspase 3/7 activity assays were performed with the Caspase-Glo 3/7 Assay System (Promega) as per the manufacturer’s instructions.

### ApoBD isolation

Following apoptosis induction by UV-irradiation, ApoBDs generated from DC2.4 cells were isolated by differential centrifugation as described previously (Atkin-Smith *et al*, 2017; Phan *et al*, 2018). Briefly, cultured supernatant were centrifuged at 300 x *g* for 5 min to sediment cells and debris. Supernatants were further centrifuged at 3,000 x *g* for 10 min to enrich ApoBDs. ApoBD purity was determined by flow cytometry and confocal microscopy as described below. Diameter of ApoBDs was quantified as per previous studies (Santavanond *et al*, 2024).

To isolate ApoBDs from BMDCs, peptide pulsed or unpulsed BMDCs were plated in 15 cm tissue culture dish in SF-RPMI/BSA and UV irradiated (150 mJ/cm^2^) to induce apoptosis. Following 4 h incubation at 37°C, apoptotic supernatant (containing both cells and ApoBDs) was collected, centrifuged at 3,000 x *g* (collecting cells and ApoBDs without pelleting small EVs) and resuspended in FACS sorting buffer (2 mM EDTA, 5% FBS in 1x PBS). Samples were stained with 1:200 A5-FITC and 0.5 µM TO-PRO-3 in 1x A5 binding buffer for 10 min followed by isolation of based on A5^high^/FSC^low^ gating using the FACS Aria III (BD Biosciences). Apoptotic supernatant was also collected for staining with 1:200 A5-APC, 0.5 µM PO-PRO-3 and 1:200 IgG-PE or H2K^b^(SIINFEKL)-PE) and analysed by flow cytometry. Data analysis was performed using FlowJo software 8.8.10 (Tree Star).

To isolate ApoBDs from IAV-infected DC2.4 cells (22 h post infection), supernatant from DC2.4 cell culture were collected and centrifuged at 3,000 *g* for 10 min. Samples were stained with A5-FITC in 1x A5 binding buffer for 10 min, centrifuged (3,000 x *g*, 10 min) and pellets were resuspended in FACS sorting buffer (1x PBS, 25 mM HEPES pH 7.0, 1 mM EDTA, 1% FBS). Samples were filtered through 70 μm cell strainers and ApoBDs (A5^high^/FSC^low^) were sorted using the FACS Aria III (BD Biosciences).

### Time-lapse DIC microscopy

For differential interference contrast (DIC) time-lapse imaging of cells undergoing apoptosis, Cells were seeded into a 4- or 8-well Nunc® Lab-Tek® II chambered coverglass slides (Thermofisher Scientific) in SF-RPMI/BSA and imaged for ∼4-6 h at 37°C with 5% CO_2_. IAV(PR8 strain)-infected DC2.4 cells were imaged for ∼16 h at 37°C with 5% CO_2._ Imaging was performed using the Zeiss Spinning Disk Confocal microscope with 63x/1.4 oil immersion objective. Images were processed using Zen Blue software (Zeiss).

### Confocal fluorescent microscopy

For live cell imaging with fluorescent probes for cell surface marker, ∼ 3×10^4^ LPS-activated or immature DC2.4 cells and ∼6×10^4^ HMDCs were seeded into an 8-well Nunc® Lab-Tek® II chambered coverglass slides (Thermofisher Scientific) in SF-RPMI/BSA. HMDCs were stained with Hoechst 33342 and primary antibodies of human Anti-HLA-DR-PE (1:10) (Miltenyi Biotech) following induction of apoptosis. In certain experiments, apoptotic DC2.4 cells were identified by A5-FITC and TO-PRO-3 or Hoechst 33342 staining. Live cell imaging was performed on either a Zeiss Confocal LSM 780 PicoQuant FILM or Zeiss Confocal LSM 800 microscope using x63 oil immersion (Zeiss).

### Transmission electron microscopy

DC2.4 cells were exposed to UV irradiation at 150 mJ/cm^2^ using a CX-2000 UV crosslinker (UVP Inc.) and incubated at 37°C, 5% CO_2_ for 2 or 4.5 h, respectively. Cells were fixed by Karnofsky solution, washed by distilled water, then fixed in an aqueous solution of 1% OsO_4_ (for membrane fixation) and treated with 1% uranyl acetate solution. Fixed cells were embedded in Epon 812, and thin sections were cut and stained with uranyl acetate and lead citrate for observation under a Jeol-1010 electron microscope (Jeol) at 80 kV.

### Flow cytometry

For apoptosis analysis by flow cytometry, cells or cell fragments were stained with 1:200 A5-FITC and 0.5 µM TO-PRO-3 in 1x A5 binding buffer for 10 min at room temperature and analysed by BD FACS Canto II flow cytometer (BD Biosciences). Flow cytometry data was analysed using FlowJo software 8.8.10 (BD Biosciences) as described previously (Jiang *et al*, 2016). For flow cytometric analysis of surface markers, cells or cell fragments were stained with primary antibodies: anti-mouse CD40-FITC, anti-mouse MHCI-Pacific Blue, anti-mouse CD80-FITC and anti-mouse CD86-FITC (BD Biosciences) at a 1:100 dilution for 30 min at 4°C in combination with A5-FITC or A5-V450 and TO-PRO-3. Isotype-matched controls were used for each primary antibody.

### Immunoblotting

Cell and ApoBD lysates were prepared using cell lysis buffer (20 mM HEPES pH 7.4, 1% IGEPAL® CA-630, 10% glycerol, 1% Triton X-100, 150 mM NaCl, 50 mM NaF, protease inhibitor cocktail tablet (Roche, CH)) and 20-40 μg of protein was separated on 4%-12% gradient SDS-PAGE gel and transferred onto nitrocellulose or PVDF membranes. Membranes were blocked with 5% skimmed milk in PBS containing 0.1% Tween® 20 (PBST) (Sigma-Aldrich) for 1 h and probed overnight at 4°C with the following primary antibodies (1:1,000 unless stated otherwise): anti-cleaved caspase 3 (Cell Signaling; D175), anti- pro caspase 3 (Santa Cruz; sc7148), anti-pannexin1 (1:500; Cell Signaling; D9MIC), anti-ROCK1 (Santa Cruz; sc17794), anti-TAP1 (1:100; Sigma-Aldrich; SAB2102370), anti-β actin (Sigma-Aldrich; AC-74), anti-ERK2 (Santa Cruz; D-2). Membranes were incubated in secondary horseradish peroxidase-conjugated sheep anti-mouse (1:5,000; Millenium Science), donkey anti-rabbit (1:5,000; Abcam) or goat anti-rabbit (1:5,000; Millenium Science) antibodies in 5% skimmed milk in PBST for 1 h. HRP signal was developed using ECL Prime (GE Lifesciences) and captured using the Syngene G:Box gel Documentation and Analysis System (Syngene).

### Antigen presentation assay (OVA system)

For antigen presentation assays, OT.I T cells were isolated from the spleen of C57BL/6 transgenic OT.I mice. All experiments were approved by the La Trobe University Animal Ethics Committee in accordance with the National Health and Medical Research Council Australia code of practice for the care and use of animals for scientific purposes. To generate peptide pulsing of BMDCs, peptides SIINFEKL_257-264_ or gBT-1_498-505_ were prepared at 10^−5^ M in SF-RPMI. BMDCs were resuspended in diluted peptide and incubated on ice for 1 h, resuspending the cells gently every 10 min to prevent cell sedimentation. Cells were then washed in SF-RPMI, induced to undergo apoptosis and ApoBDs were isolated by FACS as described in ApoBD isolation. Co-culture of BMDCs with ApoBDs was then performed: Naïve OT.I T cells were stained with 1 µM CFSE (Life Technologies) and 5×10^4^ cells per well were seeded into a 96-well plate. Sorted BMDC ApoBDs (generated from unpulsed, SIINFEKL-pulsed and gBT-1-pulsed BMDCs) were resuspended to an equal concentration and added to OT.I T cells according to indicated ratios. T cells were also cultured alone or with the SIINFEKL peptide. Cells were incubated at 37°C with 5% CO_2_ for 72 h, then samples were stained with anti-Vα2TCR-PE and analysed by flow cytometry using the BD FACS Canto II and FlowJo Software (BD Biosciences).

### Antigen presentation assay (IAV system)

To induce DC apoptosis, DC2.4 cells were infected with 10 MOI PR8 (strain A/Peurto Rico/8/1934 H1N1) for 22 h and DC-ApoBDs were collected via differential centrifugation or FACS sorting as described in ApoBD isolation. Infected DCs and DC-ApoBDs were co-cultured at a 1:2 and 1:5 ratio, respectively, with NP_366-374_-specific CD8^+^ T cells (Zanker *et al*, 2013) for 5 h in the presence of 10 μg/mL Brefeldin A (Sigma). Samples were stained with anti-CD8-APC and fixed with 1% paraformaldehyde. Samples were washed and permeabilised with 0.2% saponin and stained with anti-IFN-γ-FITC. All samples were analysed on the FACS CANTO II and using FlowJo Software (BD Biosciences).

### Statistics

Data are presented as the mean ± the standard error of the mean (SEM). Statistical analysis was performed using an unpaired Student’s two-tailed *t*-test or two-way analysis of variance (ANOVA) with Sidak’s multiple comparisons test.

## Author contributions

A.L.H., J.P.S., S.C., B.S., A.A.B. and I.K.H.P. designed the experiments with input from other co-authors, in particular W.C., S.O., M.D.H., T.K.P. and G.K.A.-S. A.L.H., A.A.B. and I.K.H.P. wrote the manuscript. All figures were assembled by A.L.H., S.C., B.S. and I.K.H.P. J.P.S., A.A.B., G.F.R. and G.K.A.-S. assisted the *in vivo* irradiation model. S.C. and S.O. performed experiments on the disassembly of apoptotic BMDCs and DC-ApoBD-mediated antigen-presentation using the OT-1 system. B.S. assisted with the isolation and characterisation of DC-ApoBDs derived from DC2.4 cells, and analysis of Ca_v_3.2 expression by immunoblotting. L.J., S.A., I.Y., and S.S. performed TEM studies. R.T. and T.K.P. assisted with the generation of TAP1^−/−^ DC2.4 cells. A.L.H. performed all other experiments.

## Declaration of interests

The authors declare no competing interests.

## Acknowledgements

The authors would like to thank the La Trobe University Bioimaging Platform and La Trobe Animal Research and Teaching Facility for their technical supports. This work was funded by the National Health and Medical Research Council (GNT1173662 to I.K.H.P.) and Australia Research Council (DP200100458 to I.K.H.P.).

---

**Supplementary Figure 1.**
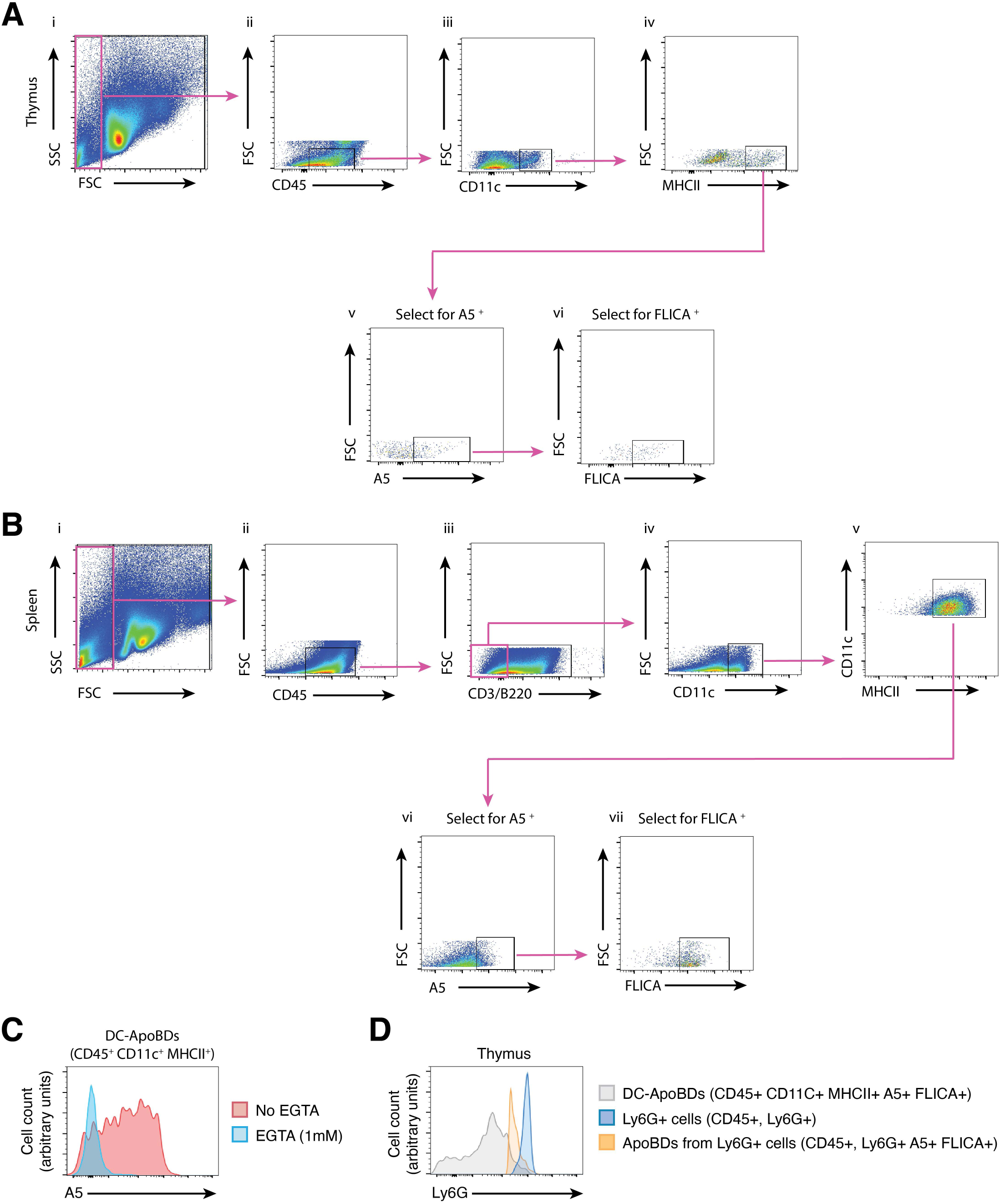
Electronic gating strategy used to identify DC-derived ApoBDs within the thymus and spleen. **(A)** Gating steps for thymic samples: (i) whole thymic events, FSC^low^ events (i.e. large EVs) were gated from other events (i.e. viable cells) and further gated upon. (ii-iv) DC-derived large EVs were selected as CD45^intermediate^, CD11c^intermediate^ and MHCII^intermediate^ events. (v-vi) A5^high^ and FLICA^high^ events were selected, determining an apoptotic origin. DC-derived ApoBDs from the thymus are defined as a FSC^low^, CD45^intermediate^, CD11c^intermediate^, MHCII^intermediate^, A5^high^ and FLICA^high^ population. **(B)** Gating steps for splenic samples: (i) whole spleen events, FSC^low^ events (i.e. large EVs) were gated from other events (i.e. viable cells) and further gated upon. (ii-v) DC-derived ApoBDs were selected as CD45^intermediate^, CD3/B220^low^, CD11c^intermediate^ and MHCII^intermediate^ events. (vi-vii) A5^high^ and FLICA^high^ events were selected, determining an apoptotic origin. DC-derived ApoBDs from the spleen are defined as a FSC^low^, CD45^intermediate^, CD3/B220^low^, CD11c^intermediate^, MHCII^intermediate^, A5^high^ and FLICA^high^ population. **(C)** A5 binding to DC-ApoBDs in the presence or absence of EGTA (1 mM), as determined by flow cytometry. **(D)** Monitoring the staining of Ly6G on DC-ApoBDs by flow cytometry, with Ly6G^+^ cells and ApoBDs as a comparison.

**Supplementary Figure 2.**
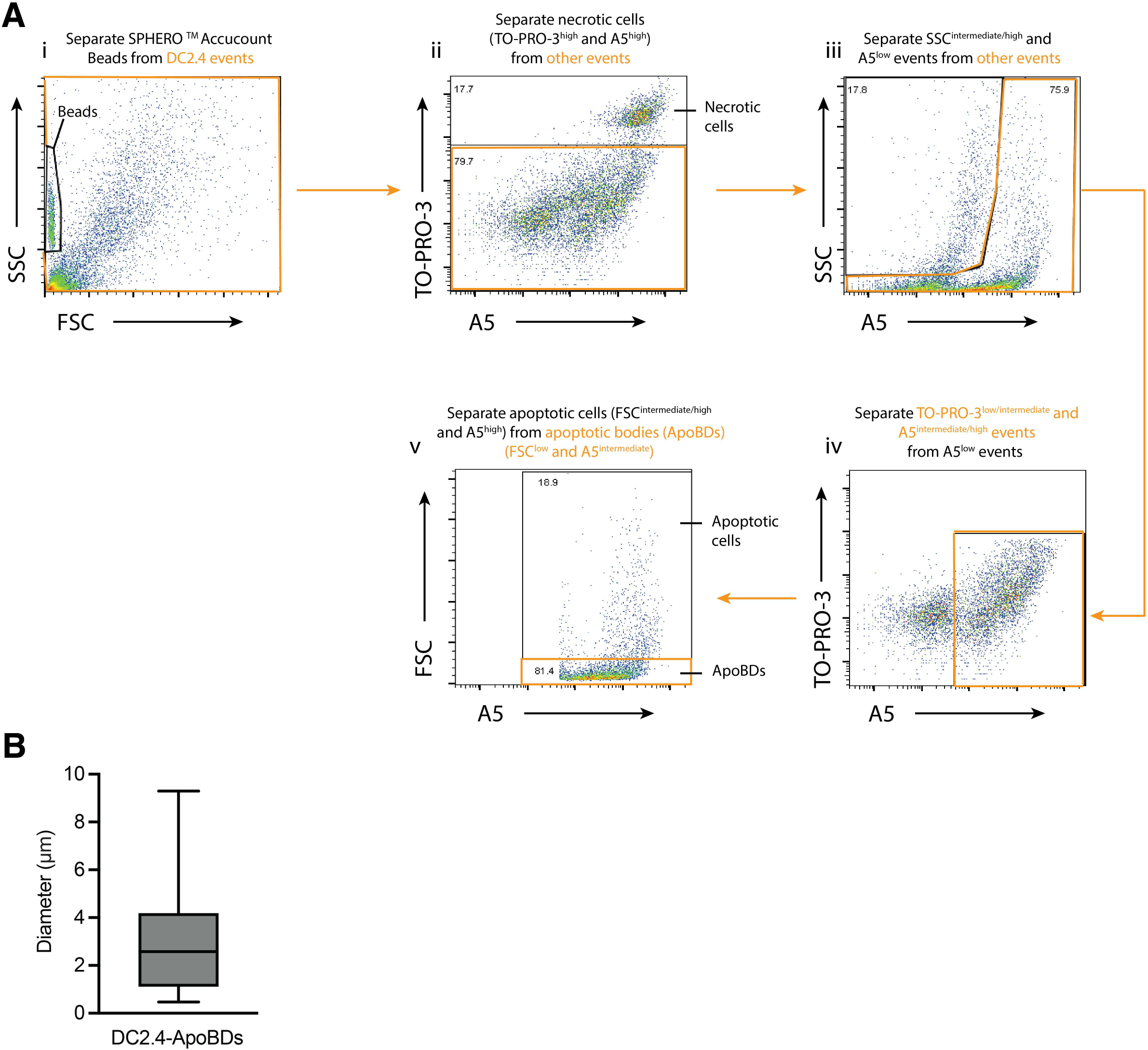
Electronic gating strategy used to identify and size quantification of DC2.4-derived ApoBDs. **(A)** Gating steps for DC2.4 cell sample: (i) DC2.4 events were separated from SPHERO AccuCount Beads and further gated upon. (ii) Necrotic cells (TO-PRO-3^high^ and A5^high^) were separated from other events (TO-PRO-3^low/intermediate^). (iii) Viable cells (SSC^intermediate/high^ and A5^low^) were separated from other events. (iv) Apoptotic events (TO-PRO-3^low/intermediate^ and A5^high^) was separated from A5^low^ events (i.e. debris). Apoptotic cells (FSC^intermediate/high^ and A5^high^) were separated from ApoBDs (FSC^low^ and A5^intermediate^). **(B)** Quantitation of live microscopy data from Figure 1L to determine the diameter of A5 positive cells/fragments.

**Supplementary Figure 3.**
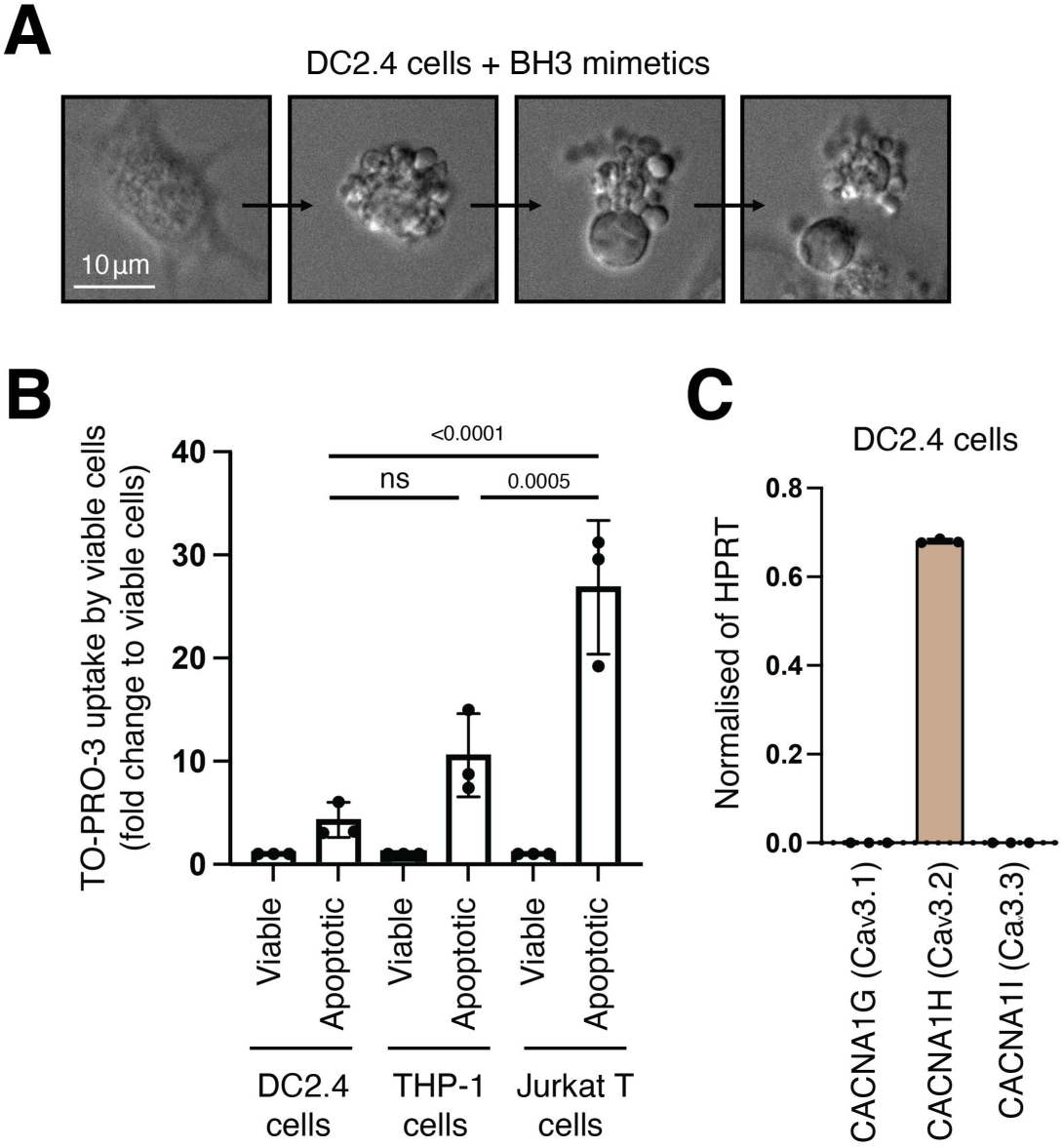
Induction of DC apoptosis by BH3 mimetics treatment. Time-lapse DIC microscopy images monitoring the disassembly of apoptotic DC2.4 cells treated with BH3 mimetics (ABT-737, 2 μM; S63845, 5 μM) to induce apoptosis.

**Supplementary Figure 4.**
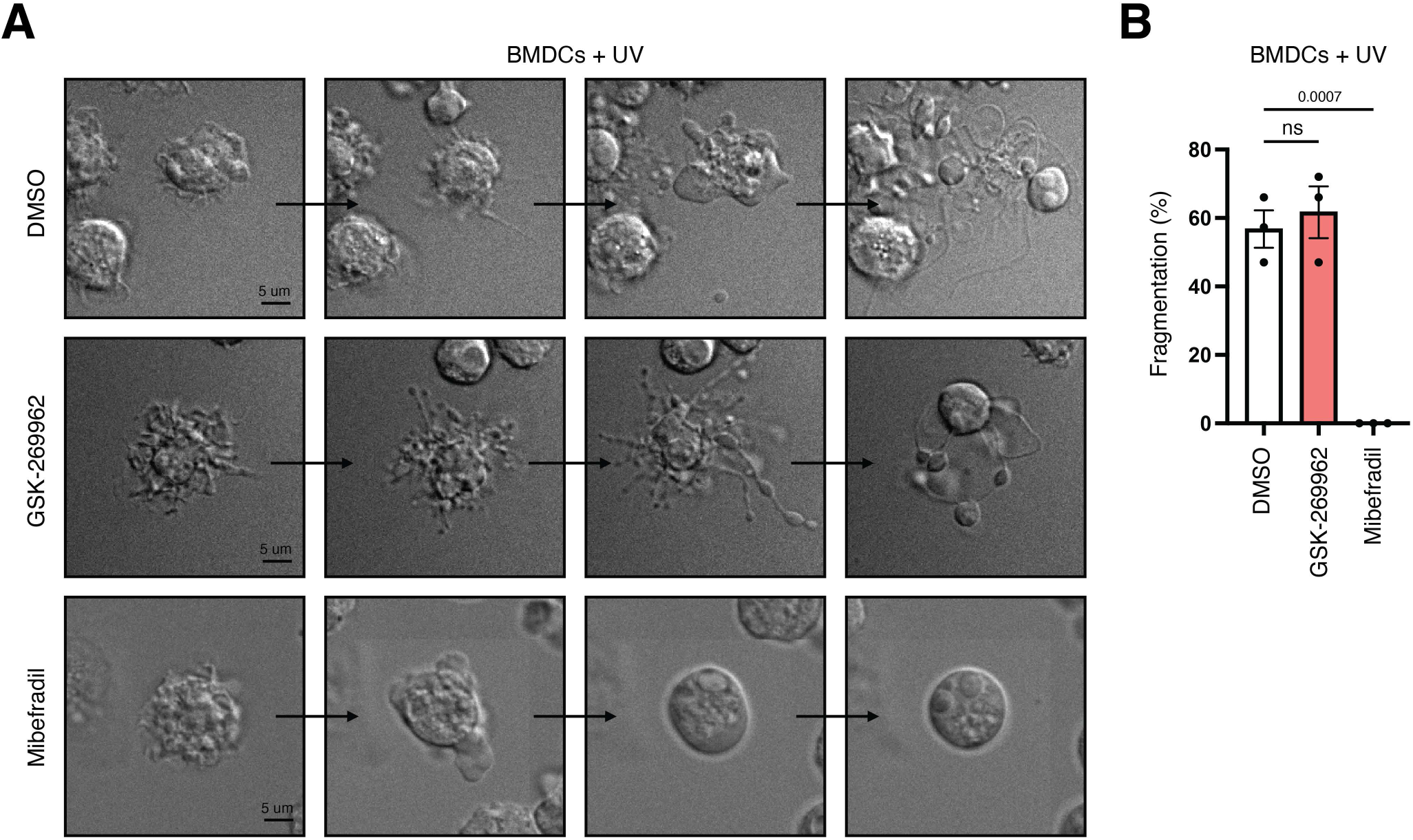
Mechanism of BMDC disassembly during apoptosis. **(A)** Time-lapse DIC microscopy images monitoring the formation of membrane blebs and apoptopodia by BMDCs treated with UV irradiation to induce apoptosis and in the presence of vehicle control (DMSO), the ROCK1 inhibitor GSK-269962 (1 μM) or the T-type calcium channel inhibitor mibefradil (10 μM). Data are representative of three independent experiments. **(B)** Quantitation of live microscopy data from (A) to determine the percentage of apoptotic BMDCs that underwent cell fragmentation (*n*=3). Error bars represent s.e.m. Statistical analysis was performed using two-way ANOVA with Sidak’s multiple comparisons test, ns= P>0.05.

**Supplementary Figure 5.**
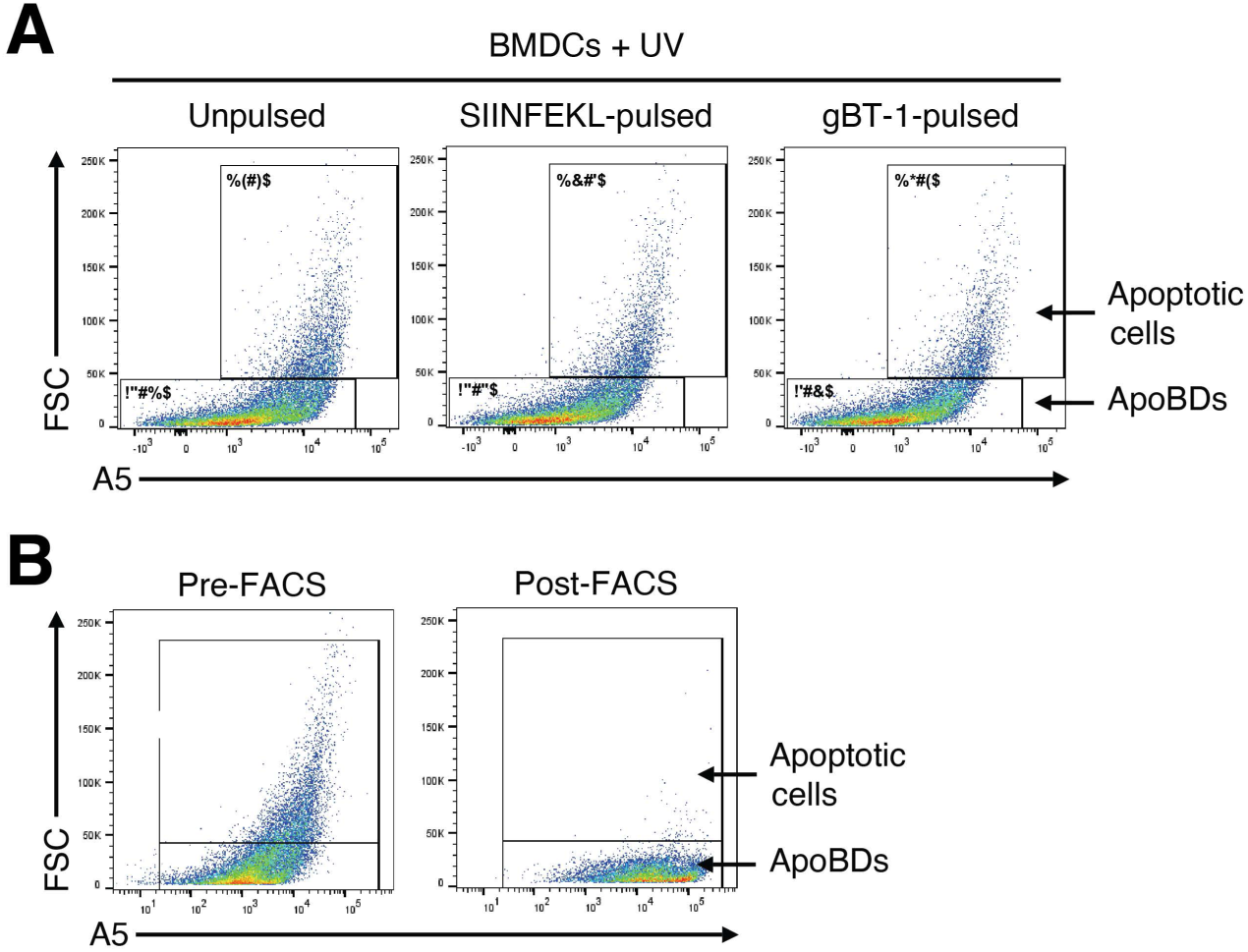
Characterisation of BMDCs cell death and ApoBD isolation. **(A)** Representative flow cytometry plots of UV-irradiated unpulsed, SIINFEKL-pulsed or gBT-I-pulsed BMDCs, gated for apoptotic cells and ApoBDs. **(B)** Representative flow cytometry analysis of the level of apoptotic cells and ApoBDs in samples pre- and post-FACS to isolate ApoBDs following UV irradiation of BMDCs.

**Supplementary Figure 6.**
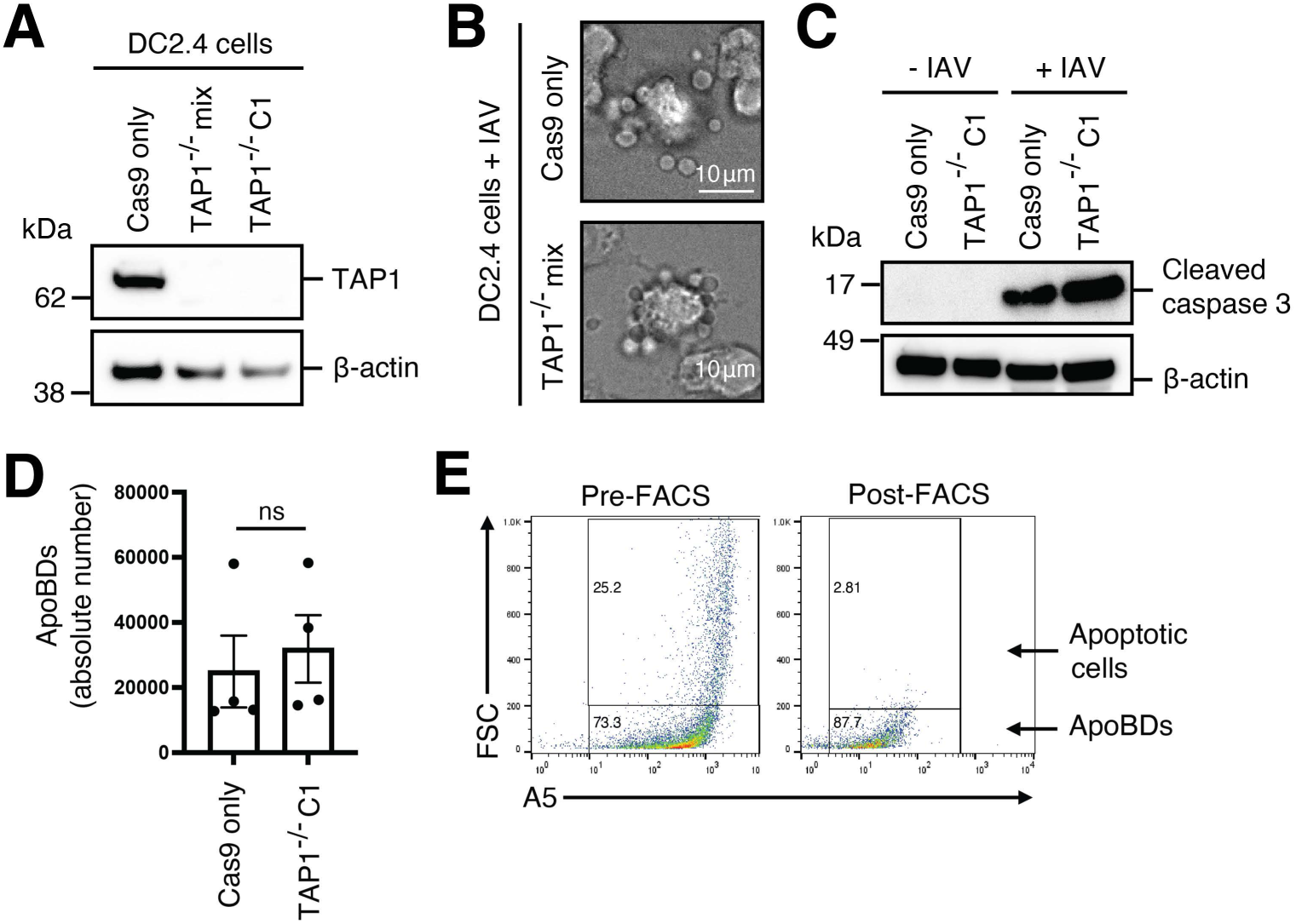
Induction of DC apoptosis by IAV infection. **(A)** TAP1 protein expression in Cas9 only, TAP1^−/−^ mix population (mix) or TAP1^−/−^ clone 1 (C1) DC2.4 cells was determined by immunoblotting. **(B)** DIC microscopy images monitoring the disassembly of apoptotic DC2.4 cells induced to undergo apoptosis by IAV infection. **(C)** Immunoblot analysis of caspase 3 activation in IAV-infected Cas9 only or TAP1^−/−^ C1 DC2.4 cells (*n*=2). **(D)** Flow cytometry analysis of ApoBDs generated from Cas9 only or TAP1^−/−^ C1 DC2.4 cells following IAV infection. 1.5×10^5^ DC2.4 cells were induced to undergo apoptosis. **(E)** Representative flow cytometry analysis of the level of apoptotic cells and ApoBDs in samples pre- and post-FACS to isolate ApoBDs following IAV infection of DC2.4 cells. Error bars represent s.e.m. Statistical analysis was performed using unpaired Student’s two-tailed *t*-test, ns= P>0.05. Data are representative of at least two independent experiments.

## Notes

### Competing Interest Statement

The authors have declared no competing interest.

